# Persistent impairment of spatial hearing and neural binaural interaction after “temporary” noise-induced hearing loss

**DOI:** 10.64898/2026.01.16.699944

**Authors:** Chase A. Mackey, Jane A. Mondul, Amy N. Conner, John Peacock, Monica A. Benson, Troy A. Hackett, Daniel J. Tollin, Ramnarayan Ramachandran

## Abstract

Many people have trouble understanding speech in noisy environments despite normal audiometric thresholds, a condition referred to as “hidden hearing loss.” A leading hypothesis attributes this deficit to inner hair cell synaptic pathology (cochlear synaptopathy), which can persist after recovery from noise-induced temporary elevation of audiometric thresholds. This pathology impairs the temporally precise sound encoding necessary for spatial hearing, but direct evidence in primates, human or nonhuman, is lacking. Here, we show that a single noise exposure, producing only a temporary threshold elevation, induced long-lasting alterations in synaptic morphology, without synapse loss, resulting in persistent spatial hearing deficits for up to 11 months post-exposure in nonhuman primates of both sexes. These perceptual deficits were accompanied by a reduction in a neurophysiological measure of binaural processing - the binaural interaction component (BIC) of the auditory brainstem response (ABR). Both behavioral and neural deficits persisted despite full recovery of audiometric thresholds. Together, these findings provide the first evidence in primates that noise exposure disrupts spatial hearing and binaural neural circuit function without loss of hair cells or synapses. Because spatial hearing tests and the ABR/BIC are clinically accessible, this work also establishes translational biomarkers for early neural dysfunction underlying hidden hearing loss.

## Introduction

Hearing in noisy environments is crucial to survival across species and is particularly important for human communication. Understanding the neural mechanisms that support speech perception in noise is a central goal in auditory neuroscience and has direct implications for diagnosing and treating hearing difficulties. Although impaired hearing in noise is commonly associated with elevated audiometric thresholds, many individuals with clinically normal thresholds also experience substantial difficulty, a phenomenon often referred to as “hidden hearing loss” (Bakay et al., 2018; Bharadwaj et al., 2015; Parthasarathy et al., 2020; Schaette and McAlpine, 2011; Tremblay et al., 2015). These difficulties are prominent in tasks requiring binaural processing of suprathreshold sound, which has the real-world implication of diminished capacity for discriminating between competing sounds in social settings (Bharadwaj et al., 2015; Cooper and Gates, 1991; Parthasarathy et al., 2020; Ruggles et al., 2011). Over the past two decades, cochlear synaptopathy, which refers to the loss or dysfunction of inner hair cell ribbon synapses, has emerged as a leading candidate mechanism underlying hidden hearing loss. Synaptopathy can be induced by noise exposure that only temporarily elevates audiometric thresholds (e.g. Kujawa and Liberman, 2009; Song et al., 2016; Valero et al., 2017). In rodent models, such exposures lead to auditory nerve fiber loss and robust neurophysiological effects, including reduced Wave I amplitude of the auditory brainstem response (ABR), degraded temporal sound envelope encoding, and binaural integration (Benson et al., 2025; Shaheen et al., 2015; Shaheen and Liberman, 2018). Of particular relevance, Benson et al. (2025) suggested the binaural interaction component (BIC) of ABR is sensitive to synaptopathy in guinea pigs. The BIC is a residual waveform resulting from subtracting the sum of the monaurally evoked ABRs from the binaurally evoked ABRs. It has been studied as a biomarker for binaural hearing abilities (Anbuhl et al., 2025; Benichoux et al., 2018) and correlates with binaural behavioral abilities in normal and hearing-impaired subjects (Laumen et al., 2016; Van Yper et al., 2016) . Although not easily measurable clinically in human populations (Sammeth et al., 2020), we recently established its basic characteristics in nonhuman primates (Peacock et al., 2021). The BIC thus provides a reliable means to assess the impact of noise exposure and synaptopathy on neural and behavioral measures of spatial hearing.

Despite compelling evidence from rodent models, described above, identifying reliable behavioral or neurophysiological assays for synaptopathy in humans has proven difficult. Human studies have yielded mixed results, with some finding indirect evidence linking putative synaptopathy to deficits in binaural processing and speech-in-noise perception (Bharadwaj et al., 2015; Bramhall et al., 2020; Parthasarathy et al., 2020; Ruggles et al., 2011), including confirmation from human temporal bone studies where cochlear histopathology is directly assessed (Wu et al., 2021, 2020, 2019). Other studies, however, have failed to detect clear associations between suspected synaptopathy and functional outcomes (Bramhall et al., 2019; Guest et al., 2018). Differences in stimulus design and analytic approaches likely contribute to these discrepancies (Bramhall et al., 2019; DiNino et al., 2021), highlighting the need for translational models that link cochlear pathology, neural processing, and perception within the same subjects.

To address this gap, we recently developed a nonhuman primate (NHP) model of noise-induced hearing loss that bridges rodent and human studies. The model was developed by utilizing different levels of noise exposure (Valero et al., 2017), and within-subject correlations between cochlear histopathology, brainstem physiology, and behavior (Burton et al., 2020; Hauser et al., 2018; C. A. Mackey et al., 2021), in a species with close anatomical, physiological, and perceptual similarity to humans (Burton et al., 2019; Hackett et al., 2001; Hoglen et al., 2018; Joris et al., 2011; C. Mackey et al., 2021; Moody, 1994; Stebbins, 1982). Importantly, NHPs exhibit reduced susceptibility to noise-induced cochlear damage relative to rodents (Burton et al., 2020, 2019; Hauser et al., 2018; Hawkins et al., 1976; Valero et al., 2017). Our recent cochlear histological work demonstrated that a single 120 dB SPL exposure to octave band noise for 4 hours produces a temporary threshold shift and persistent inner hair cell synaptic hypertrophy without loss of synapses or hair cells (Mondul et al., 2025). Although the functional consequences of these synaptic morphological changes remain unknown, synaptic hypertrophy has been linked to glutamatergic excitotoxic mechanisms (Pal et al., 2025; Robertson, 1983; Sheets et al., 2017). Moreover, cochlear histopathology is dynamic after noise exposure (Benson et al., 2025; Hickman et al., 2021, 2020; Kim et al., 2019; Liu et al., 2012; Shi et al., 2013; Wu et al., 2024), and synaptic recovery has been proposed to underlie improvements in binaural neural processing and behavior ∼2 months after exposure (Benson et al. 2025). Motivated by these findings, the present study tests the hypothesis that noise exposure causing only temporary threshold shifts leads to persistent alterations in inner hair cell synapses, binaural brainstem processing, and behavioral measures of spatial hearing.

## Materials and methods

### Subjects

Ten adult rhesus macaques (*Macaca mulatta*) ages 5 – 10 years old with body weights ranging from 5 – 13 kg were used in this study. Three male and four female macaques comprised the behavior group (n = 7), while 12 macaques (4 female) comprised the neurophysiology group. Macaques were fed a commercial diet (LabDiet Monkey Diet 5037 and 5050, Purina, St Louis, MO) supplemented with fresh produce and foraging items. Macaques were also provided manipulanda as well as auditory, visual, and olfactory enrichment on a rotational basis. Macaques were fluid-restricted for the study and received filtered municipal water, averaging at least 20 ml/kg of body weight/day (typically closer to 25 ml/kg/day). Their weight was monitored at least weekly (typically 4–5 days per week) and remained within the reference weights set to index the animal’s health while on study. Macaques were maintained on a 12:12-h light:dark cycle, and all procedures occurred between 8 AM and 6 PM during their light cycle. After repeated behavioral assessments to identify compatible social partners, these macaques were individually housed (due to incompatibility with available cohorts for social housing) with visual, auditory, and olfactory contact with conspecifics maintained within the housing room. The housing room was in an AAALAC-accredited facility in accordance with the *Guide for the Care and Use of Laboratory Animals,* the Public Health Service Policy on Humane Care and Use of Laboratory Animals, and the Animal Welfare Act and Regulations. Macaques in this colony received routine health assessments and tuberculosis testing twice yearly. All research procedures were part of protocols that were approved by the Institutional Animal Care and Use Committee at Vanderbilt University Medical Center (VUMC). We have previously published on the sound levels in the vivarium, which were found to be in a safe range (McLeod et al., 2022). Spatial behavioral experiments were performed after extensive audiometric and physiological characterization (ABRs, DPOAEs), confirming normal hearing.

### Surgical procedures

Monkeys were prepared for chronic experiments using standard techniques employed in nonhuman primate studies, and as reported in previous studies (Dylla et al., 2013; Mackey et al., 2023). Briefly, anesthesia was induced via administration of ketamine and midazolam and maintained via isoflurane. A titanium headpost was implanted on the skull to restrict head movement during head fixation, minimizing changes in sound pressure level at the ear during positioning across behavioral sessions. The headpost was secured to bone using 8mm titanium screws (Veterinary Orthopedic Instruments) and encapsulated in bone cement (Zimmer Biomet). Multimodal analgesics (pre- and post-procedure), intra-procedure fluids, and antibiotics (intra-procedure) were administered to the monkeys under veterinary oversight. Further details about these procedures can be found in Dylla et al. (2013).

### ABR and DPOAE Threshold Testing

We have previously published on our clinical metrics of hearing threshold (e.g. Stahl et al. 2022). A Scout biologic system (Natus) was used to measure distortion product otoacoustic emissions (DPOAEs). DPOAEs were elicited using pairs of tones (f1 and f2; f2 > f1) delivered via an in-ear probe assembly. Primary tone frequency ratios and level relationships were chosen to evoke robust emissions across the 0.5-10 kHz range. To estimate DPOAE thresholds, emission amplitudes were recorded across a series of decreasing stimulus levels. The distortion product was considered present when its amplitude exceeded both an absolute level criterion (0 dB SPL) and the surrounding noise floor by 6 dB. Threshold was defined as the lowest stimulus level at which a repeatable emission meeting these criteria could be detected.

Auditory brainstem response (ABR) threshold was measured and calculated consistent with our previous methods (e.g., Stahl et al. 2022). We utilized tone bursts (0.5-32 kHz) presented at 27.7 Hz, with frequency dependent rise/fall times: 1ms (2-32 kHz), 2ms (1 kHz), or 4ms (0.5 kHz); plateau durations: 0.5 ms (2-32 kHz), 1ms (1 kHz) or 2 ms (0.5 kHz)). We measured the ABR using subdermal needle electrodes (Rhythmlink) in a vertex-to-mastoid montage, with a ground electrode on the shoulder. Electrode impedances were ≤ 3 kOhm.

### Behavioral Apparatus and Stimuli

The apparatus used for these behavioral experiments has been described in previous publications (C. A. Mackey et al., 2021; Rocchi et al., 2017). Briefly, monkeys were seated in a custom-designed primate chair situated inside a sound-treated booth (IAC, model 1200A and Acoustic Systems ER 247). Sounds were delivered from a free-field speaker (Rhyme Acoustics NuScale 216) located in the frontal field at 90 cm from the center of the monkey’s head. For the spatial experiments, speakers were placed at different angles on the azimuthal plane. Signals and noise were presented from different speakers for the 22.5, 45, 67.5, and 90-degree conditions of the spatial release of masking experiments. The speakers were calibrated with a ¼ inch microphone (378C01, PCB Piezotronics) that was located just outside the monkey’s ear canal. Care was taken to ensure that the speaker outputs were within 3 dB across all frequencies to avoid speaker-specific cues. Calibrations were also performed at the different speaker locations used in the spatial experiment to ensure that the sound level at the head was as specified. Experimental flow was controlled by a computer running OpenEx software (System 3, TDT Inc., Alachua, FL). Tones and noise were generated using a sampling rate of 97.6 kHz. The lever state was sampled at a rate of 24.4 kHz, yielding a temporal resolution of ∼ 40 μs for the lever release.

### Binaural Interaction Component Apparatus and Stimuli

Methods used for recording the binaural interaction component (BIC) were recently reported by the authors (Peacock et al., 2021), and mirror previous publications by the Tollin lab (Benichoux et al., 2018; Sammeth et al., 2020), with the exception that the data presented here were recorded while animals were under gas isoflurane anesthesia (1.5-2%). Briefly, steel subdermal needle electrodes (Grass Technologies) were placed at the apex along the interaural axis between the ears, with a reference electrode at the nape of the neck, and a ground electrode on the monkey’s shoulder. Stimuli were generated and evoked potentials were recorded via an RME Fireface UCX sound card (RME audio) and a World Precision Instruments (WPI) ISO-80 biological amplifier. Custom eartips were used to deliver click stimuli with variable interaural time differences via MATLAB. Clicks were 90 dB SPL presented at an average rate of 33 Hz via Tucker Davis Technology MF-1 speakers, which were calibrated with a Bruel and Kjær type 4182 microphone. ITDs were -1000 to 1000 µs in 500 µs steps. Conditions were interleaved and presented 3000 times each.

### Task Structure

Monkeys performed a lever-based reaction-time Go/No-Go tone-detection task. Details of the task have been reported in our previous publications (Dylla et al., 2013; C. A. Mackey et al., 2021; Rocchi et al., 2017). Briefly, monkeys initiated a trial by pulling a lever. Trials could be signal trials (80%) in which a tone signal of fixed duration was played after a random delay period of 800-3500 ms after lever was pulled, or they could be catch trials (20%), in which no tone signal was played. The monkey was required to release the lever within a response window (600 ms after tone onset) to indicate detection on signal trials and to continue holding the lever on catch trials. Lever release on signal trials (hits) were rewarded with fluid. Lack of lever release within 600 ms of the onset of the tone on signal trials (misses) was taken to indicate non-detection and was not rewarded or punished. Lever release on catch trials (false alarm) resulted in a 6-10 s timeout in which no trial could be initiated. Correct rejections (lack of release on catch trials) were not rewarded. Experiments were blocked by tone frequency and masker location, while tone level was varied trial by trial, using the method of constant stimuli.

### Noise exposure

Monkeys were treated with atropine and anesthetized with a mixture of ketamine (10 mg/kg) and midazolam (0.05 mg/kg). They were then intubated, following which anesthesia was maintained with isoflurane (1.5 – 2%). The exposure noise was delivered through a closed-field acoustic system. MF1 speakers (TDT Inc.) coupled with probe tips were inserted into the ear canal. The stimulus was an octave-band (2000-4000 Hz) of noise presented for four hours at 120 dB SPL to both ears, which elicits temporary threshold shifts in macaques (Mondul et al. 2025). We verified that the noise level varied by < 1 dB over the exposure time window.

### Behavioral data analysis

Signal detection theoretic methods were used to analyze behavioral performance, following previous studies (C. Mackey et al., 2021; Macmillan and Creelman, 2004; Tanner and Swets, 1954). Behavioral performance from each block of data was analyzed to calculate hit rate at each tone level (*H(level)*) and false alarm rate (*F*). From these values, the discriminability of signals was calculated as 𝑑^’^(𝑙𝑒𝑣𝑒𝑙) = (𝑧(𝐻(𝑙𝑒𝑣𝑒𝑙)) − 𝑧(𝐹)-, where *z* represents the conversion to a standard normal variate, and was implemented via the function “norminv” in MATLAB. From discriminability, the behavioral accuracy was given by the probability correct, *pc(level)*, as 𝑝𝑐(𝑙𝑒𝑣𝑒𝑙) = 𝑧^-1^(𝑑^’^(𝑙𝑒𝑣𝑒𝑙)⁄2), where *z^-1^*converted standard normal variates into behavioral accuracy. The psychometric function was defined as the relationship between *pc(level)* and the sound pressure level (SPL) of the tone. The psychometric function was fit with a modified Weibull cumulative distribution function (CDF) 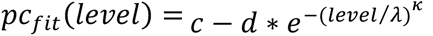 𝑓𝑜𝑟 𝑙𝑒𝑣𝑒𝑙 ≥ 0, where *c* represents saturation and *d* represents the range of the function, and *λ* and κ represent the threshold and slope parameters, respectively. The threshold was calculated as the tone level at which *pc_fit_(level)* = 0.76. The dynamic range of the function was calculated as the x values spanned from 0.1*saturation to 0.9*saturation, as in our previous work (C. Mackey et al., 2021). Reaction times were calculated for all hit trials as follows:

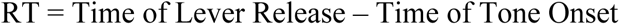

### BIC Analysis

The BIC is calculated by averaging the ABR signal in each condition with filter cutoffs of 30 Hz and 3 kHz. The BIC was calculated as the difference between the ABR amplitude evoked in the binaural stimulation condition and the algebraic sum of the monaural conditions, with the left and right ear monaural ABRs shifted by ITD/2 to compensate for the delay in the ITD conditions before summing them together (see Peacock et al., 2021).

### Euthanasia and Cochlear Histological Analysis

Comprehensive cochlear histological analyses of the nonhuman primate subjects in this study were recently published (Mondul et al., 2025). We provide a brief overview here. Following completion of the current study, NHPs were euthanized using sodium pentobarbital and sodium phenytoin (Euthasol; >120 mg/kg IV) and transcardially perfused with 0.9% phosphate-buffered saline and 4% phosphate-buffered paraformaldehyde (PFA). Cochlear tissue was harvested for dissection and immunohistochemistry, the details of which are extensively described in Mondul et al. (2025). Immunolabeling and confocal imaging of cochlear whole mounts were conducted to quantify IHC and OHC counts, IHC and OHC ribbon counts and sizes, and efferent terminal densities (Burton et al., 2020; Mondul et al., 2025; Valero et al., 2017). Data from noise-exposed subjects were compared to roughly age-matched unexposed subjects to assess anatomical integrity along the cochlear length.

### Statistical analysis

In all cases, curve fits were obtained using the non-linear least-squares method implemented in MATLAB. Statistical effects reported were assessed by using linear mixed effects models (“fitlme”) in MATLAB. This allowed us to accommodate datasets with variable numbers of subjects and conditions, a common reason for avoiding repeated measures ANOVA (Krueger and Tian, 2004). P-values were obtained by likelihood ratio testing of the model with the effect in question against the model without the effect in question.

## Results

Spatial hearing was evaluated in a cohort of seven nonhuman primates (Rhesus macaques) before and after exposure to a four hour, 120 dB SPL octave band noise. A detailed analysis of audiologic testing and cochlear histopathology associated with this noise exposure has recently been published (Mondul et al., 2025) and is summarized here (Figure 1). The temporary threshold shift was verified using distortion product otoacoustic emissions (DPOAEs) and ABR thresholds (Figure 1A, Figure 1B). Histology revealed little to no hair cell loss, little to no hair cell synapse loss, and robust hair cell synaptic hypertrophy at the 2 and 10 month time-points (Figure 1C).

**FIGURE 1.**
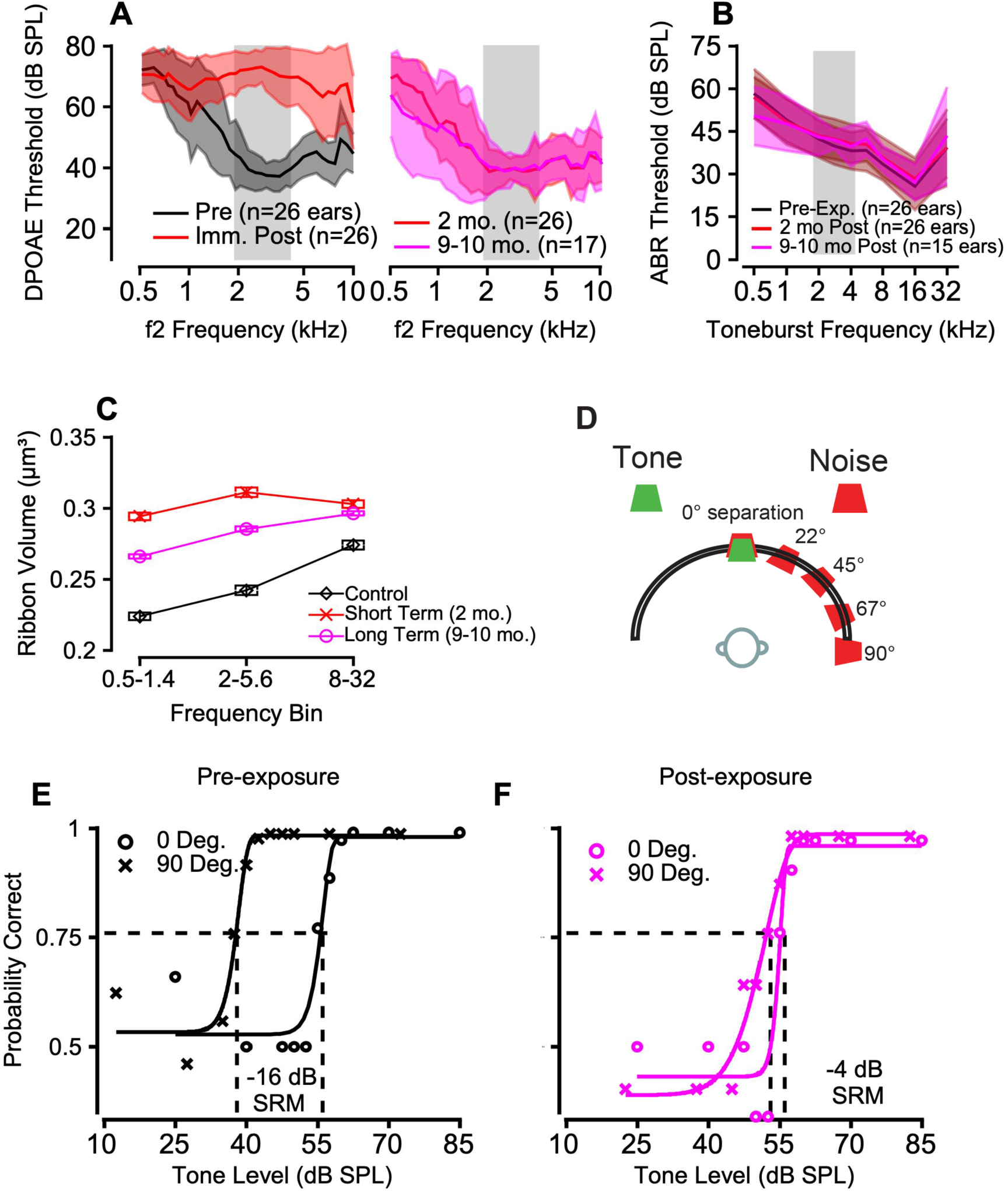
Effects of temporary threshold shift-inducing noise exposure on spatial release of masking in male nonhuman primates. **A**) Distortion product otoacoustic emissions (DPOAE) thresholds before (black), immediately post-, and at subsequent time-points following noise exposure confirmed the temporary depression of outer hair function. **B)** Auditory brainstem response threshold (the lowest sound level evoking a detectable ABR) confirmed the temporary nature of the threshold shift pre (black) and at subsequent time-points post-noise exposure. **C)** Inner hair cell ribbon volumes (Mean +/-SEM) were significantly larger in noise exposed NHPs compared to controls. **D**) Apparatus used to measure spatial hearing in nonhuman primates. Tones and broadband, continuous noise were presented from a loudspeaker. Noise could be co-localized with the tone (the 0 degree condition), or spatially separated, which elicits spatial release of masking, or **SRM**. **E**) Representative psychometric functions from monkey Bi when the noise was colocalized with the 5.656 kHz tone (circles), and spatially separated (x-hairs). SRM is the difference between detection thresholds in the two conditions. **F**) Psychometric functions from Monkey Bi 10 months after noise exposure indicating reduced SRM.

### Spatial release of masking (SRM)

After the noise-induced temporary threshold shift (TTS) had resolved (Figure 1A, Figure 1B), SRM was measured at 22.5, 45, 67.5, and 90 degrees (Figure 1D), at an early (∼2 months) and a late (∼10 months) time-point to address the dynamic nature of spatial processing after noise exposure (see Introduction). Representative psychometric functions showing how SRM was calculated, as the difference between detection thresholds, are shown in Figure 1E. SRM was reduced compared to pre-exposure values following noise exposure, due to large elevations in threshold in the spatially separated conditions (Figure 1F). SRM of 5.6 kHz tones was impaired across speaker locations in the three male monkeys (Figure 2A). Noise exposure also impaired SRM in a cohort of female macaques (Figure 2B; Table 1). A mixed effects model confirmed the effect was significant (“SRM ∼ Noise Location + Tone Frequency*Noise Exposure Status + (1|Monkey)”; interaction between noise exposure and tone frequency: t = 4.11, df = 192, p = 5*10^-5^). SRM was reassessed 10-11 months post noise exposure, and still indicated a deficit (pink data points, Figure 2A-B). Female macaques displayed greater variability than male macaques in that the frequency specificity of SRM deficits changed over time. For example, Monkey Lu exhibited an initial deficit at 5.6 kHz, consistent with male macaques, but at 2.8 kHz a deficit emerged at 9 months that was not present at the 3-month time-point. In contrast, Monkey Pi displayed no deficit at 5.6 kHz at 3 months, but at 9 months, SRM decreased at several spatial separations.

**FIGURE 2.**
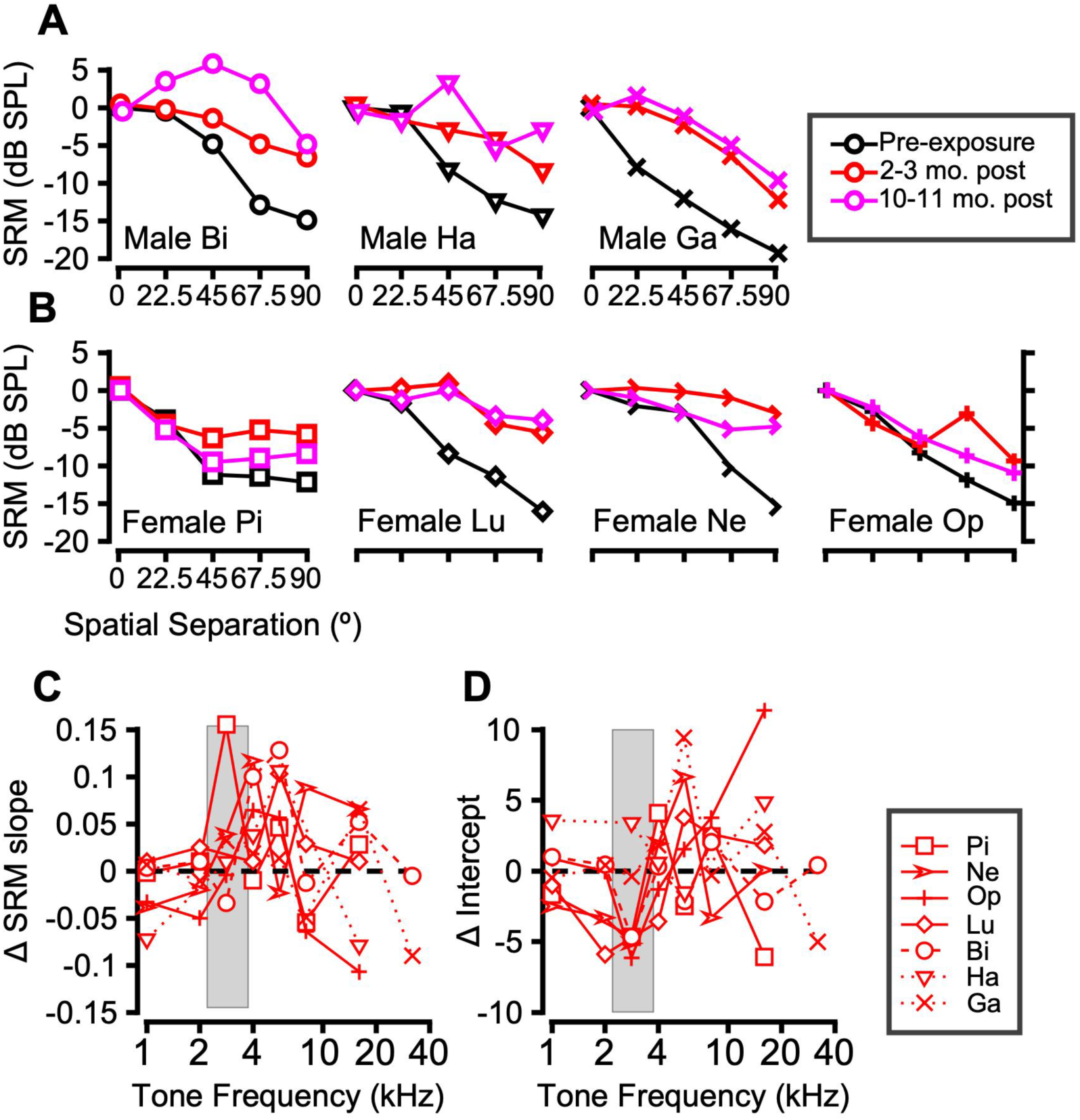
Effect of noise exposure on spatial hearing in nonhuman primates (*n =* 7). **A)** Spatial release of masking (SRM) of 5.6 kHz tones is shown as a function of spatial separation for each male macaque monkey at the pre exposure (black), 2-3 month post (red), and 10-11 month post time-points (pink). **B)** SRM is shown as a function of spatial separation for each female macaque monkey at the pre exposure (black), 2-3 month post (red), and 9-10 month post time-points (pink). Tone frequencies shown are as follows: 2.8 kHz (Monkey Pi), 5.6 kHz (Lu), 4 kHz (Ne), and 5.6 kHz (Op). **C)** Change in the slope of the SRM vs. spatial separation trends as a function of all tone frequencies used in the study for each macaque at the early time-point. **D)** Change in the intercept of the SRM vs. spatial separation trends as a function of tone frequency for each macaque at the early time-point. A grey box indicates the frequency content of the noise-exposure band.

**Table 1.**
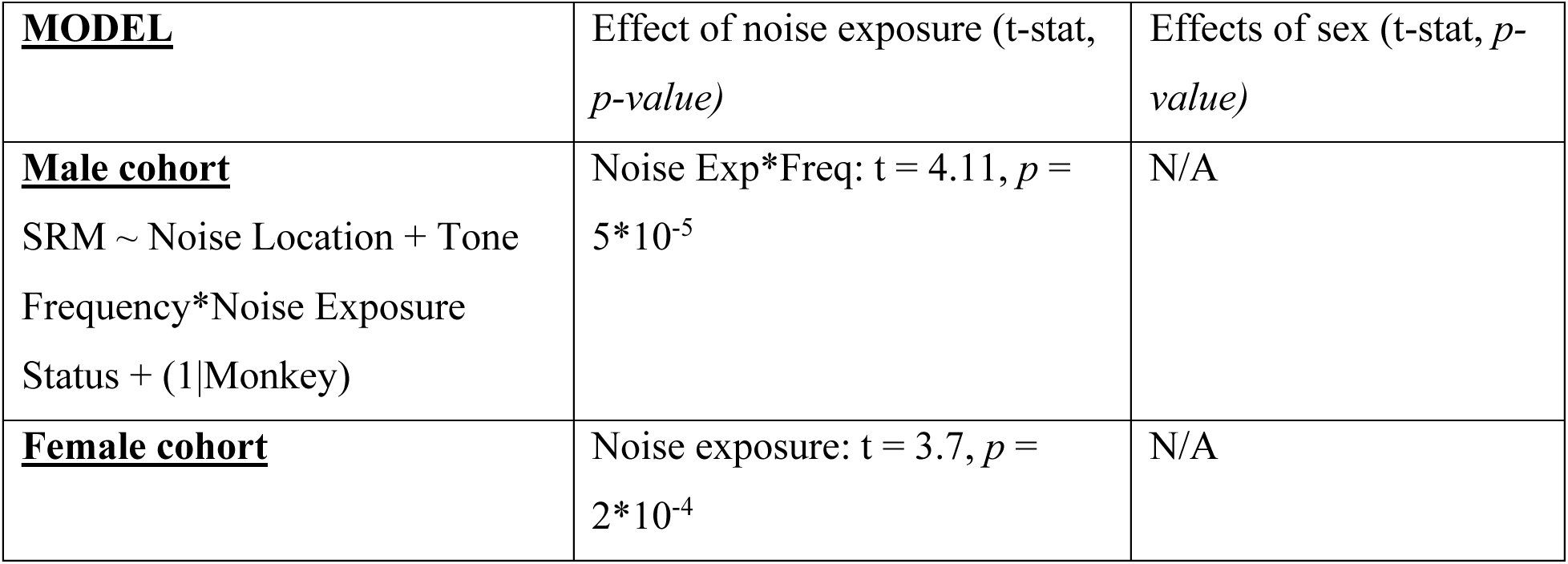
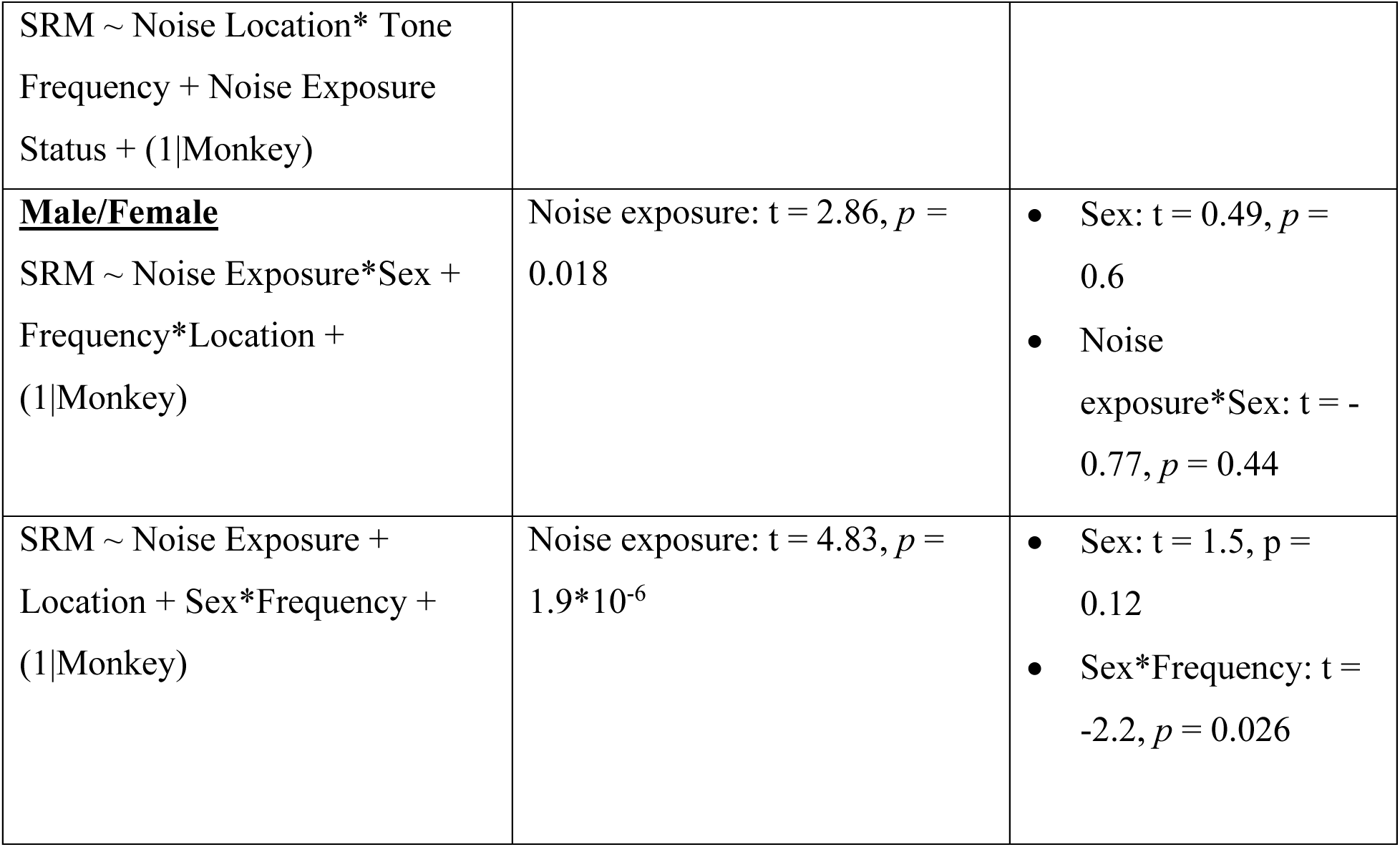
Mixed effects model analysis of the effects of noise exposure and sex on SRM. Note: effects of sex are descriptive and should be interpreted with caution due to the small sample size of the study (n=3 male, n=4 female).

| **<u>MODEL</u>** | Effect of noise exposure (t-stat, *p-value*) | Effects of sex (t-stat, *p-value*) |
| --- | --- | --- |
| **<u>Male cohort</u>**<br>SRM ∼ Noise Location + Tone<br>Frequency*Noise Exposure<br>Status + (1\|Monkey) | Noise Exp*Freq: $t = 4.11$, $p = 5 \times 10^{-5}$ | N/A |
| **<u>Female cohort</u>** | Noise exposure: $t = 3.7$, $p = 2 \times 10^{-4}$ | N/A |
| SRM ~ Noise Location* Tone<br>Frequency + Noise Exposure<br>Status + (1\|Monkey) |  |  |
| **<u>Male/Female</u>**<br>SRM ~ Noise Exposure*Sex +<br>Frequency*Location +<br>(1\|Monkey) | Noise exposure: $t = 2.86, p = 0.018$ | • Sex: $t = 0.49, p = 0.6$<br>• Noise exposure*Sex: $t = -0.77, p = 0.44$ |
| SRM ~ Noise Exposure +<br>Location + Sex*Frequency +<br>(1\|Monkey) | Noise exposure: $t = 4.83, p = 1.9 \times 10^{-6}$ | • Sex: $t = 1.5, p = 0.12$<br>• Sex*Frequency: $t = -2.2, p = 0.026$ |

SRM values at different spatial separations were fit with lines to estimate slope and intercept, providing a qualitative summary measure of SRM that could be displayed across different tone frequencies (Figure 2C-D). This accompanies the mixed effects model analysis that quantifies these observations. Deficits due to noise exposure could manifest as a decrease in slope or an increase in intercept. SRM slope for 5.6 kHz tones decreased after noise exposure for monkeys Bi and Ha, while SRM intercept at 5.6 kHz increased in Monkey Ga. All seven monkeys displayed slope reductions at least one frequency in the 2.8-5.6 kHz range.

We conducted an analysis of potential differences between the male and female cohorts that is merely descriptive, due to the small sample size in our study (*n =* 3 male, *n =* 4 female). The similarities and differences in noise induced deficit between the sexes was described by a mixed effects models (Table 1) that took the form “SRM ∼ Noise Exposure*Sex + Frequency*Location + (1|Monkey)” (Effect of sex: t = 0.49, df = 405, p = 0.6; interaction of noise-exposure and sex: t = -0.77, df = 405, p = 0.4). The observation that noise-induced deficits occurred at a broader range of frequencies in female macaques (2.8-5.6 kHz) was quantified by the significant Sex by Frequency Interaction (Table 1). The testing of interactions between sex and noise exposure is crucially different from simply comparing the effects of noise exposure between cohorts, which is a common mistake when investigating sex differences (Garcia-Sifuentes and Maney, 2021). A significant interaction indicates that sex modifies the effect of noise exposure, whereas merely comparing the main effect of noise separately in the male and female cohorts does not directly test for an effect of sex. The effect of noise exposure on SRM in the female cohort was significant, as with the male cohort (t = 3.7, df = 209, p = 0.0002), and the effect of noise exposure on SRM in the dataset with both sexes was significant (p = 1.9*10^-6^; Table 1). It should be noted that NHP studies with small sample sizes are not ideal for evaluating sex differences, and these results should be interpreted cautiously (see Discussion).

### Reaction times and psychometric function slope

Guided by the hypothesis that noise exposure would affect other measures of accuracy and/or speed, a goal of the present work was to characterize its effects on the slope of the psychometric function and on reaction times (RTs). Previous work has documented how various stimulus features affect the slope of the psychometric function (C. Mackey et al., 2021; Rocchi and Ramachandran, 2020), which can be quantified by the psychometric dynamic range (DR). DR was defined as the tone levels spanned by the sloping portion of the Weibull curve fit (see Methods and (C. Mackey et al., 2021)). DR slightly increased following noise exposure, as shown in Figures 3A and 3C. This effect was small but significant across the dataset in a mixed effects model (Coefficient estimate: 1 dB; t = 2.6, df = 405, p = 0.01). The effect was restricted to the most spatially separated conditions (67.5 and 90 degrees), coinciding with the greatest changes in accuracy. Sex differences were not significant, and sex did not interact significantly with noise exposure in the mixed-effects model of DR (*t = 0.9, df = 405, p = 0.3)*.

**Figure 3.**
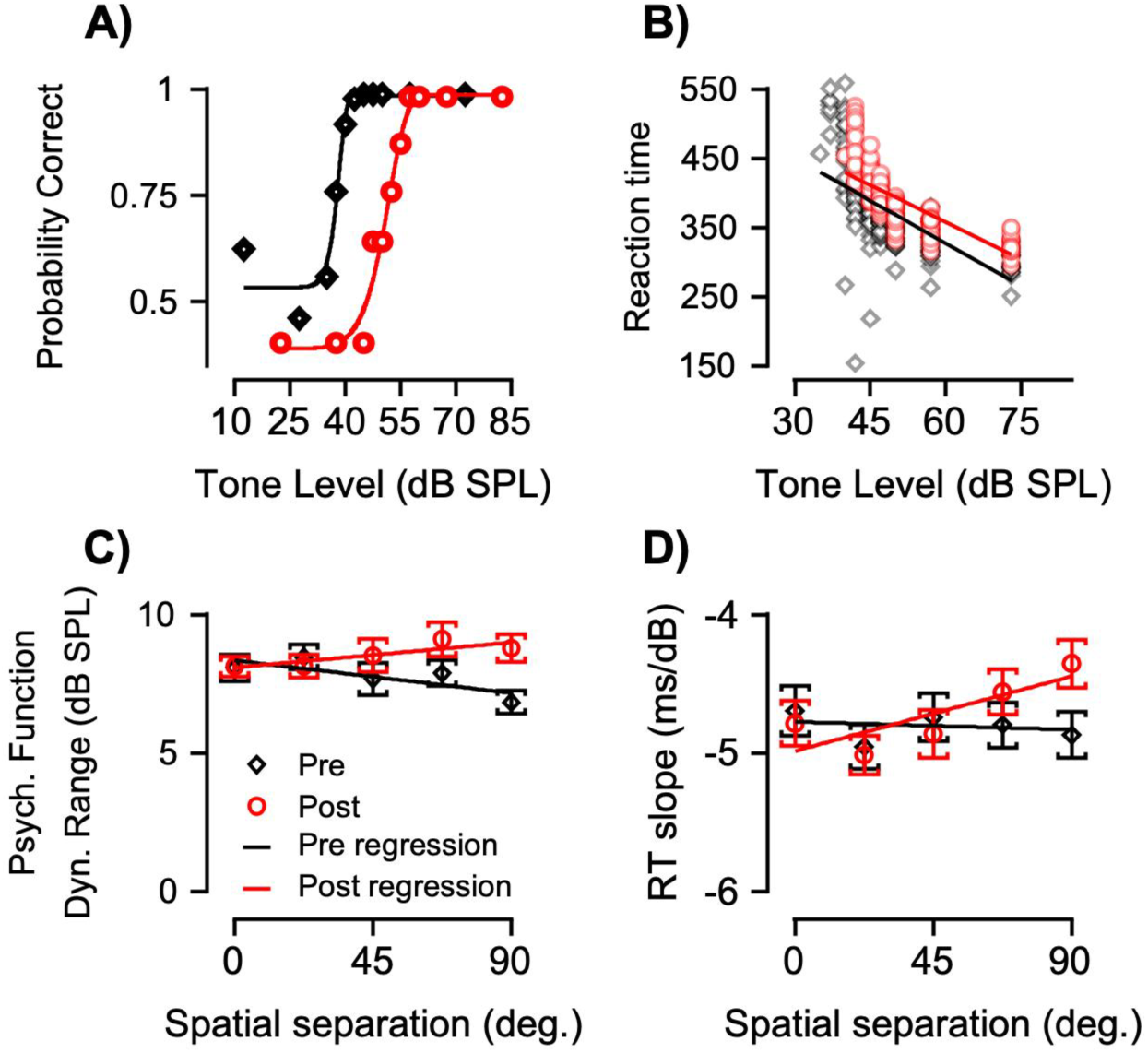
Effects of noise exposure on psychometric function dynamic range and reaction times. **A**) Representative psychometric functions from monkey Bi pre- and post-noise exposure. **B**) Representative reaction times from monkey Bi pre- and post-noise exposure. **C**) Mean psychometric function dynamic range across the dataset (*n =* 405 psychometric functions). Lines represent the linear regression in a multifactorial mixed-effects model. **D**) Mean reaction-time slopes across the dataset (*n =* 405 slopes). Error bars represent the standard error of the mean. In all panels, black and grey indicate pre-exposure data, red indicates post-exposure data.

Previous work has characterized how RTs change with sound level, and signal-to-noise ratio (Dylla et al., 2013; Kemp, 1984; Kemp and Irwin, 1979). Namely, RTs decrease as sound level increases, and conversely RTs increase as SNR decreases. These effects were quantified using a linear fit to RT as a function of tone level (Figure 3B). RT slope changed after noise exposure, which could be modeled as an interaction between noise location and noise exposure status (t = 2.29, df = 405, p = 0.023). The interaction indicates that before noise exposure RT slope did not change as a function of noise location, but following noise exposure RT slope decreased as a function of location. This decrease indicates a reduction in the facilitatory effects of tone level on detection speed. The biggest effect across the dataset was, on average, 0.4 ms/dB decrease in RT slope at 90 degrees, which amounts to a 12 ms average change in RT. Sex differences were not significant, and sex did not interact significantly with noise exposure in the mixed-effects model of RT slope or intercept (*p = 0.10, p = 0.19,* respectively).

### Correlation with cochlear histology

Having characterized perceptual changes in spatial processing following noise exposure that alters inner hair cell ribbon morphology, it was of interest to assess the extent to which frequency-specific cochlear damage predicts frequency-specific spatial hearing deficits. Two measures of interest emerged from our analysis of the cochlear histology: ribbon synapse counts and ribbon synapse volume (Mondul et al., 2025). We correlated change in SRM, relative to pre-exposure values, with relative synapse per IHC at the corresponding cochlear frequency place (normalized to the 1 kHz place; a measure that has been found to adequately account for individual variability in synapse counts, (Mondul et al. 2025). Correlations were not significant for twelve out of thirteen ears from seven NHPs (Pearson correlation, *p > 0.05*). Similarly, frequency-specific correlations between changes in SRM and ribbon synapse volume relative to a control group were not significant in 11 of 13 ears from 7 NHPs.

## Binaural interaction after noise exposure

The finding that noise exposure impairs spatial hearing suggests the encoding of localization cues is compromised. This positions the binaural interaction component (BIC) of the ABR as a promising noninvasive measure of cochlear synaptic integrity (see Introduction). To capitalize on this opportunity, the BIC of the ABR was measured in male and female macaques before and after noise exposure. The BIC is a derived measure of the click-evoked ABR, calculated by subtracting the algebraic sum of monaural ABRs from the binaural ABR. Example evoked potentials and calculation of the BIC are shown in Figure 4. Figure 4 A and B show monaural ABRs in response to clicks, which are summed to give the dashed trace in Figure 4C. The BIC is calculated as the difference between the summed trace and the binaural trace (evoked response to simultaneous clicks to both ears, solid trace, Figure 4C). Figure 4D shows the BIC traces, exhibiting a stereotypical negativity at approximately 4 milliseconds, which, after the sign is inverted, will be referred to as the BIC amplitude. Figure 4D shows monkey Pi’s BIC traces before (black traces) and after noise exposure (red traces).

**Figure 4.**
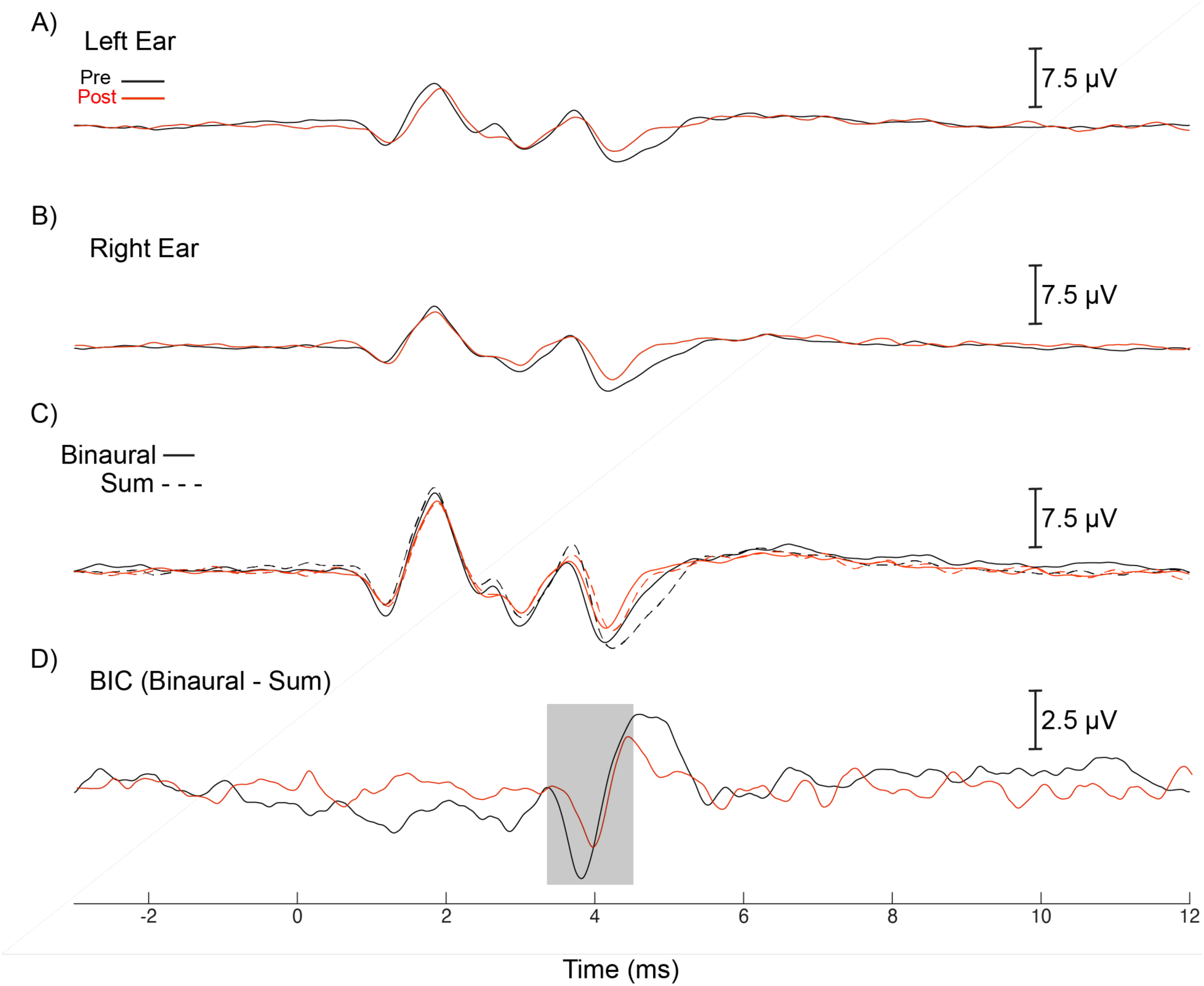
Example binaural interaction component of the auditory brainstem response from monkey Pi before (black) and after (red) noise exposure. **A, B.** Monaural ABRs in response to clicks presented to the left (**A**) and right (**B**) ears. **C.** Monaural sum (dashed) and binaural response (solid) used for the calculation of the BIC. **D.** BIC traces, with a shaded region outlining the negativity near 4 ms, or BIC amplitude.

Noise exposure reduced BIC amplitude. This near halving of the response was common, as shown in the group data displayed in Figure 5. BIC amplitudes at zero ITD are plotted for each monkey individually in Figure 5A, and BIC latencies at 0 ITD are plotted in 5B. All BIC amplitudes are plotted as a function of interaural time difference (ITD) in Figure 5C. Data were statistically analyzed using a mixed-effects model. The best-fitting model, as defined by the lowest Bayesian information criterion value, took the form *BIC ∼ ITD + Noise Exposure + ITD^2+^ ITD^2^*Noise Exposure + (1|Monkey)*. Noise exposure caused a significant reduction in BIC amplitude (*t = -9.02, df = 77, p = 1*10^-13^)*, and interacted with ITD (*t = 2.48, df = 77, p = 0.015)*. The positive t-statistic indicates that noise exposure reduced the effect of ITD (i.e., BIC vs. ITD trends were shallower following noise exposure). To visualize this statistical analysis, linear regressions taking the same form as the mixed effects model are plotted in Figure 5C, demonstrating that the 95% confidence intervals of the groups (black and red error bands) do not overlap, and that at a group level, the effect of ITD was reduced post noise exposure. There were no main effects of sex or interactions of sex with ITD or noise exposure. BIC latencies also showed no effects of sex, noise exposure, or interactions between sex and noise exposure.

**Figure 5.**
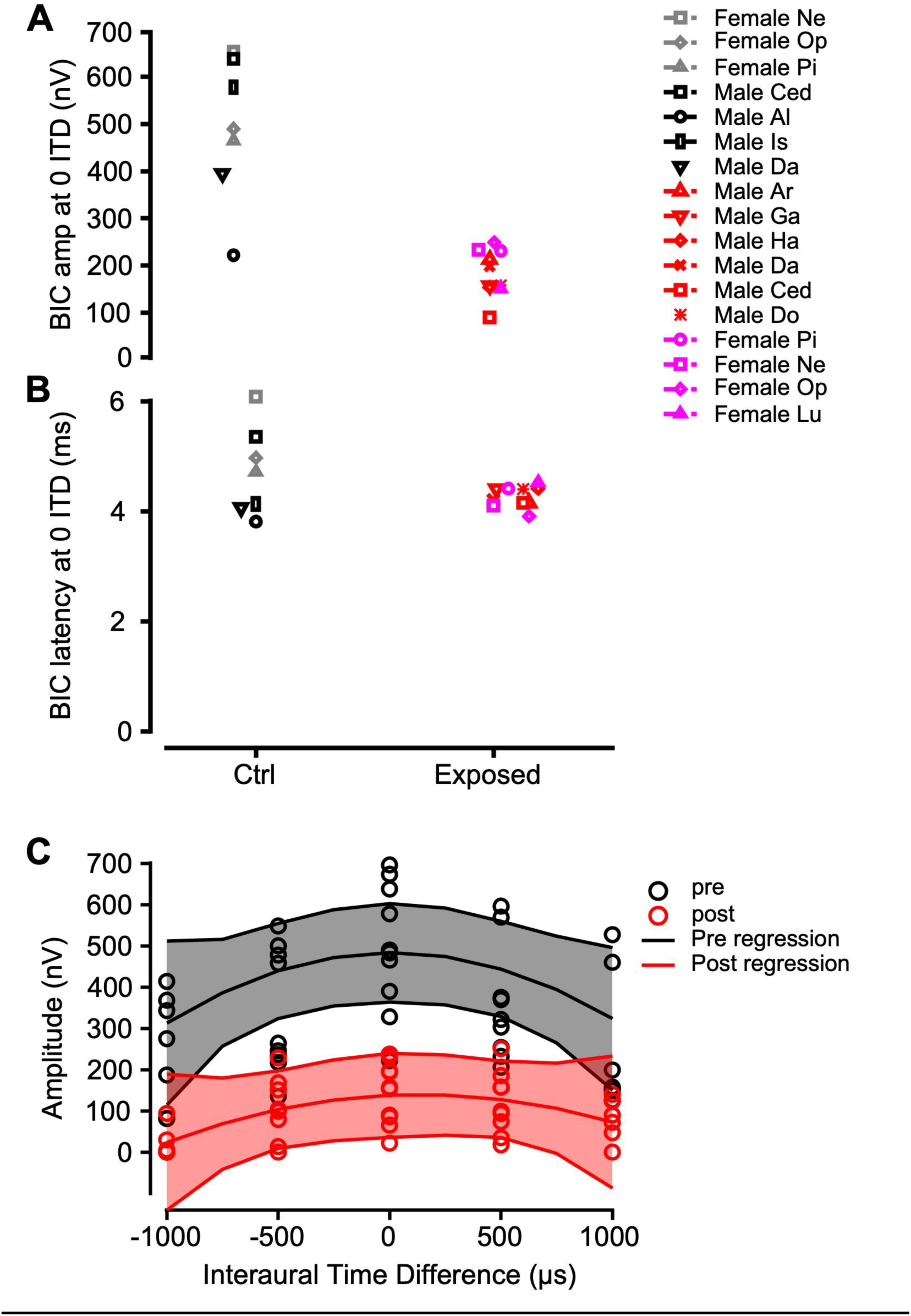
Group BIC data. **A.** BIC amplitudes of individual monkeys shown as different symbols before (black and grey) and post (red and pink) noise exposure. Male data are shown in black (pre) and red (post). Female data are shown in grey (pre) and pink (post). **B.** BIC latencies shown in the same format as A. **C.** BIC amplitudes plotted as a function of interaural time difference pre (black) and post (red) noise exposure. Data are shown as circles. Lines are multivariate linear regressions of the form *BIC ∼ ITD + Noise Exposure + ITD^2^ ^+^ ITD^2^*Noise Exposure.* The shaded error-bands show the 95% confidence intervals around the regression.

A key consideration is that the BIC is derived from Wave IV of the ABR, and Wave IV was reduced by noise exposure (Figure 4A; Figure 4B). These reductions were significant across the dataset (*Wave IV amp ∼ Noise Exposure + L/R Ear + (1|Monkey*); Effect of noise exposure, *p = 0.0013).* To quantify the degree to which BIC reductions were expected from these changes to the monaural responses, we measured a normalized BIC amplitude as *BIC_norm_ = BIC/(Wave IV_Left_ + Wave IV_Right_)*. BIC_norm_ was 0.65 (SD: 0.32) on average in the normal-hearing group, and 0.29 (SD: 0.11) in the noise-exposed group. In a mixed effects model that controlled for individual monkey differences with a random effect (BIC_norm_ ∼ Noise Exposure + (1|Monkey)), the effect of noise exposure was significant (*t =* -3.79, df = 15, *p = 0.0029*), indicating that BIC reductions exceeded those predicted by monaural scaling alone. These results mirror prior results in normal hearing guinea pigs in a test-retest experiment showing that while there could be large day-to-day fluctuations in the amplitude of the BIC, these fluctuations scaled with the amplitudes of the monaural ABRs - likely due to subtle changes in electrode impedance and placement day-to-day - but that the relative BIC amplitude remained nearly constant over time (Ferber et al., 2016). We also address monaural ABR changes in a separate publication (Conner et al. *in prep*). Here, we focus on relating binaural neurophysiology to behavioral measures of spatial hearing.

An analysis of the correlation between maximum BIC reduction and maximum SRM reduction did not yield significant results (Spearman correlation, *p = 0.53*). Similarly, BIC reduction did not correlate with individual monkeys’ increase in IHC ribbon volume (Spearman correlation, *p >* 0.4 for left and right ears’). However, this was likely partially due to the sample size, as there were only six monkeys with both pre- and post-exposure SRM and BIC data, and there is no apparent way to correlate different conditions (e.g., location vs. ITD, tone frequency, etc.). However, a strength of this dataset is that every monkey with SRM deficits also exhibited BIC reduction. Taken together, this suggests that this noise exposure, which does not elevate audiometric threshold and preferentially targets cochlear synapses, degrades temporal precision in the binaural brainstem circuitry involved in spatial hearing.

## DISCUSSION

For the first time in primates, these results demonstrate robust neurophysiological and perceptual consequences of noise exposure in the absence of detectable changes in standard clinical measures of hearing, including audiometric thresholds. Alterations in inner hair cell morphology, binaural interaction within the auditory brainstem, and spatial hearing persisted for more than ten months following the exposure. Here, we place these findings in the context of prior work on cochlear synaptopathy and “hidden hearing loss,” while acknowledging that much remains to be discovered about the relationship between the two, particularly in nonhuman primates (Mondul et al., 2025).

### Relevance to previous spatial and binaural hearing studies

While many studies have described neurophysiological changes associated with temporary threshold shifts (TTS), and others have characterized spatial hearing deficits in humans with clinically normal hearing, none have demonstrated persistent spatial processing deficits in animals with detailed cochlear histopathology. Together with perceptual studies in rodents (Benson et al., 2025; Tziridis et al., 2021) and emerging evidence in nonhuman primates (Conner et al., *in prep*; Mondul et al., 2025), our results indicate that noise exposures producing TTS can have lasting neurophysiological and behavioral consequences. In mice and guinea pigs with synaptopathy, reduced contralateral suppression in the midbrain and reduced binaural interaction in the ABR have been reported (Asokan et al., 2018; Benson et al., 2025; Shaheen and Liberman, 2018). While these effects are broadly consistent with the spatial hearing deficits and BIC changes observed here, differences in cochlear histopathology in our NHPs warrant caution in direct comparison. Our SRM results are also consistent with previous predictions from human studies of putative synaptopathy or “hidden hearing loss” (Bernstein and Trahiotis, 2020; Bharadwaj et al., 2015; Patro et al., 2021; Ruggles et al., 2011). Extending beyond threshold-based measures, we identify deficits in reaction time and psychometric function dynamic range, indicating impairments in both processing speed and perceptual sensitivity.

### Potential sensorineural mechanisms

Because the functional consequences of inner hair cell synaptic hypertrophy are still largely unknown, essential questions remain regarding how our SRM and BIC results relate to cochlear histopathology in NHPs. Unlike prior studies focused on early manifestations of cochlear histopathology following TTS, our NHPs exhibited pronounced synaptic hypertrophy through late post-exposure time points in the absence of hair cell or synapse loss (Mondul et al. 2025; our Figure 1). Synaptic hypertrophy has been linked to glutamatergic excitotoxicity and altered release dynamics, with ultrastructural evidence of reduced mitochondrial content, swollen endoplasmic reticulum, and longer response latency preceding overt synapse loss (Pal et al., 2025; Sheets et al., 2017; Robertson, 1983). Recent work further indicates heterogeneous effects across afferent populations, reflecting excitotoxic vulnerability in some fibers and altered glutamatergic signaling in surviving synapses (Moverman et al., 2023; Reijntjes et al., 2026; Suthakar and Liberman, 2022; Vincent et al., 2024). In addition, auditory nerve (AN) demyelination may contribute to post-noise perceptual and neurophysiological deficits by degrading conduction velocity and temporal precision (Kohrman et al., 2020; Wan and Corfas, 2017; Zhang et al., 2025).

Although synaptic hypertrophy has been associated with reduced AN onset jitter (Suthakar and Liberman, 2021), we nevertheless observe deficits in neurophysiological and behavioral measures that depend on precise temporal processing. This apparent discrepancy may be resolved by recognizing that binaural interaction relies on millisecond-scale temporal integration windows that are slower than auditory nerve onset responses (Brown and Tollin, 2016). Biasing AN encoding toward brief broadband transients may therefore preserve onset precision while degrading the sustained representation of spectral modulation required for binaural decorrelation in noise. Similarly, our SRM task likely depends on the sustained encoding of complex spectral cues introduced by the head and pinnae during free-field listening (Gilkey and Good, 1995). Because these spectral modulations are broad relative to tonal stimuli (Spezio et al., 2000), this may explain the lack of frequency-specific correspondence between SRM deficits and cochlear histopathology (C. A. Mackey et al., 2021). Degraded binaural integration, potentially exacerbated by AN demyelination, would be expected to propagate to the binaural brainstem. Consistent with this framework, the lateral superior olive (LSO), a likely neural generator of the BIC component of the ABR (Benichoux et al., 2018; Tolnai and Klump, 2019; Peacock et al., 2021), encodes interaural level differences (ILDs) and envelope-based timing cues critical for spatial hearing (Tollin, 2003). Impaired spectral and temporal encoding would therefore be expected to reduce both SRM and BIC amplitude, consistent with the concurrent behavioral and electrophysiological deficits observed here.

### Sex differences

We observed a modest but consistent sex difference, with female macaques exhibiting deficits across a broader range of tone frequencies than males, despite similar overall magnitudes SRM impairment and BIC reduction. In a TTS model, Benson et al. (2025) likewise reported sex-dependent effects in guinea pigs, with females showing greater susceptibility to noise-induced ABR changes but preserved spatial hearing sensitivity. Together, these findings suggest that sex may influence vulnerability or compensation following noise exposure. However, given the limited sample sizes typical of NHP studies, these observations should be interpreted cautiously. Nonetheless, they highlight the potential value of the NHP model for testing hypotheses related to sex differences in noise-induced auditory dysfunction, which are well-documented in humans (Lauer and Schrode, 2017; Pearson et al., 1995; Villavisanis et al., 2020).

### Consequences for diagnostic and therapeutic work in humans

These findings have direct implications for the development of improved diagnosis and therapeutic approaches to noise-induced hearing loss. The NHP model provides a translational platform for identifying biomarkers and candidate therapeutic targets that can be meaningfully extended to human clinical populations. Rapid clinical measures of SRM are already available (Jakien et al., 2017), and the BIC of the ABR exhibits similar characteristics in humans and macaques (Delb et al., 2003; Kelly-Ballweber and Dobie, 1984; Peacock et al., 2021; Sammeth et al., 2020). Linking behavioral and electrophysiological measures such as these with mechanistic insight from primate models may facilitate the development of more sensitive diagnostics and targeted interventions along the auditory pathway. Ultimately, such advances may help reduce the substantial global burdens of noise-induced hearing loss (Davis and Hoffman, 2019; World Health Organization, 2018).

## Acknowledgements

The authors would like to acknowledge Bruce and Roger Williams for fabrication of hardware, and Mary Feurtado for assistance with surgical procedures. The authors thank Drs. M. Charles Liberman and Leslie Liberman for conceptual input and assistance collecting histological data. The authors thank Dr. Jessica Feller, Namrata Temghare, Alejandro Tarabillo, Catherine Alek, and Jackson Mayfield for assistance collecting behavioral data. The study was supported by research grant NIH R01 DC 015988 (MPIs R. Ramachandran and B. Shinn-Cunningham). CM was supported by The Department of Hearing and Speech Sciences, and the Ruth Kirchstein pre-doctoral fellowship from NIDCD (F31 DC 019823). JAM was supported by a Ruth Kirchstein postdoctoral fellowship for audiologists from NIDCD (F32 DC 019817). DJT, JAP, and MAB were supported by NIDCD T32 DC012280 and R01 DC023100.

## Author Contributions

RR and DJT designed experiments. CAM, JP, RR and DJT conceptualized analysis. CAM, JAM, JP, MAB, and TAH collected data. CAM, JP, TAH analyzed data. CAM wrote the paper. All authors revised the paper.

The authors declare no conflicts of interest.

